# Exosomal TAR DNA binding protein 43 profile in canine model of amyotrophic lateral sclerosis: A preliminary study in developing blood-based biomarker for neurodegenerative diseases

**DOI:** 10.1101/2021.06.17.448876

**Authors:** Penelope Pfeiffer, Joan R. Coates, Yajaira M. Esqueda, Andrew Kennedy, Kyleigh Getchell, Myra McLenon, Edina Kosa, Abdulbaki Agbas

## Abstract

**Objective:** Blood-based biomarkers provide a crucial information in progress of neurodegenerative diseases with minimally invasive sampling method. Validated blood-based biomarker application in people with amyotrophic lateral sclerosis would derive numerous benefits. Canine degenerative myelopathy is a naturally occurring animal disease model to study the biology of human amyotrophic lateral sclerosis. Serum derived exosomes are potential carriers for cell-specific cargoes making them ideal venue to study biomarkers for a variety of diseases and biological processes. This study assessed the exosomal proteins that may be assigned as surrogate biomarker that may reflect biochemical changes in central nervous system.

**Methods:** Exosomes were isolated from canine serum using commercial exosome isolation reagents. Exosomes target proteins contents were analysed by Western blotting method.

**Results:** The profiles of potential biomarker candidates in spinal cord homogenate and that of serum-derived exosomes were found elevated in dogs with degenerative myelopathy as compare to control subjects.

**Conclusions:** Serum-derived exosomal biomolecules can serve as surrogate biomarkers in neuro degenerative diseases.

**Key Messages:**

- A canine with degenerative myelopathy can serve as a model animal to study human amyotrophic lateral sclerosis.
- Serum-derived exosomes contains Transactive Response DNA Binding Protein 43 (TDP-43), potential biomarker candidate.
- The levels of spinal cord TDP-43 proteins and that of serum-derived exosomes exhibited a similar profiling. Therefore, serum derived exosomes may be used as a venue for establishing blood-based biomarkers for neurodegenerative diseases.

## 1. Introduction

The aberrant protein aggregation in motor neurons is the hallmark of amyotrophic lateral sclerosis (ALS). ALS is a progressive disease that directly affects motor neurons, leading to loss of muscle function. About 5-10% of the inherited forms of ALS are linked to a mutation in the SOD1 gene (*SOD1*). The misfolded mutant [Cu/Zn] superoxide dismutase (SOD1) protein is believed to contribute to development of ALS, yet the role of misfolded non-mutant SOD1 in the disease progress is unclear. Recently, chemically modified aberrant Transactive Response DNA Binding Protein 43 (TDP-43) species were found to represent a major accumulating protein in neuronal cytoplasmic inclusions (1, 2) and in exosomes (3, 4) in Frontotemporal Lobar Degeneration (FTLD), and in ALS patients (5). Exosomes are nano-size membranous vesicles that contains several macromolecules including aberrant pathological proteins (6). The size and membranous structure of exosomes allow them to pass through the blood brain barrier (7–9). These features make exosomes a potential platform in which targeted biomolecules can be analysed.

The biomolecular link and clinical similarities between human ALS and canine degenerative myelopathy (DM) suggested that human ALS hallmark signature proteins (i.e., SOD1 and TDP-43) have the same or similar profile in canine DM. These similarities may be employed to further biomarker validity studies such as reproducibility, specificity, sensitivity, and robustness. As a naturally occur disease in the pet population, canine DM has many similarities to human ALS and is thought to be a result, in part, to mutated SOD1 protein in dogs. Thus, dogs affected with DM can serve as an animal disease model for ALS based on CNS size and complexity, and disease characteristics.

ALS is an incurable and fatal disease with a patients survival rate of 3-5 years once diagnosed (10). In most patients, the symptoms begin in the lower limbs (11, 12). Patients often complain of tripping, stumbling, foot drop, or a “slapping” gait. Upper limb involvement includes decreased dexterity in the fingers, cramping, stiffness, weakness and wrist drop (13). Early diagnosis of ALS is very difficult, which can delay time to treatment (14). Establishing a blood-based biomarker(s) will help clinicians with early diagnosis and initiating timely treatment. The pharmaceutical intervention with well-timing may improve the quality of life and lifespan for ALS patients.

ALS researchers have used rodents as an animal model with mutated SOD1 to study the pathology of the disease (15). The rodent model is of limited use as disease progression is very short (3-6 months) and is based on overexpression of the human gene for mutant SOD1. In addition, peripheral biomarker studies are difficult to perform in a longitudinal course due to limited access to serial samples of serum/exosomes in mice due to the small volume of blood. Canine DM is a late adult neurodegenerative disease accompanied by (*SOD1*) mutations (*SOD1:c.118A, SOD1:c52T*), and protein aggregation (16, 17). Early clinical signs include general proprioceptive ataxia, and spastic upper motor neuron paresis in the pelvic limbs with progression to flaccid tetraplegia and dysphagia (18–20). DM and ALS are characterized by progressive multisystem neurodegeneration involving upper and lower motor neurons. Canine DM phenotypically resembles upper motor onset ALS in reference to clinical and histopathologic features. The distribution of lesions and clinical disease progression in DM are similar to that reported for the UMN-onset ALS (21) with UMN signs in DM-affected dogs progressing later to LMN signs (16, 20).

An abnormal accumulation of TDP-43 has been demonstrated in the cytoplasm and platelets from ALS patients (22). The post translationally modified derivatives of TDP-43 (i.e., hyper-phosphorylation and ubiquitination) have been studied in ALS (23, 24). Prior research has established that the accumulation of TDP-43 in the cytosol was causal in the loss of function in affected neurons (25, 26). However, the percent of TDP-43 in peripheral blood, which represents the central nervous system (CNS) origin of TDP-43, is unknown. Therefore, assessment of the confined TDP-43 in exosomes may reflect the changes in TDP-43 protein better than that of large volume of serum or plasma where the target proteins are diluted. This study explored the feasibility of using exosomes to assess potential biomarkers for DM such as TDP-43, pTDP-43, and SOD1.

## 2. Materials and Methods

### 2.1. Animals

Previously −80^0^C frozen spinal cord tissues from thoracic region and serum samples were obtained from Dr. Joan R. Coates at the College of Veterinary Medicine, University of Missouri-Columbia under the approval of institutional animal care and use committee (IACUC# 10077). Pet owners signed an informed consent for samples being used for research purposes.

Whole blood was collected from the jugular or cephalic veins. Blood was centrifuged and serum was placed in 500 microliter aliquot tubes. Aliquots were stored at −80^0^C. The dog was necropsied for CNS tissue collections. The muscle was removed from the dorsal column of the spine. The dorsal lamina just medial to the articular processes was removed to expose the spinal cord. Specifically, the thoracic spinal cord was removed and sectioned accordingly. The tissues were immediately frozen on dry ice and stored at −80^0^C.

Tissues were harvested from various breeds with diagnosed DM and age-matched dogs not affected with DM (Table 1 in Supplemental Data). All DM affected dogs were homozygous for the SOD1 E40K mutation (20).

### 2.2. Isolation of exosomes from canine serum

The isolation of exosomes from serum of DM and control dogs was performed using a commercially available particle precipitation method following manufacturer’s protocol (miRCURY Exosome Serum/Plasma Kit, Qiagen #76603). The exosome isolation procedure was based on the capture of water molecules, which otherwise would form the hydrate envelope of particles in suspension. Mixing the starting sample with a proprietary precipitation buffer diminished the hydration of the subcellular particles and allowed the precipitation of even particles smaller than 100 nanometer with a low-speed centrifugation step (27). Before proceeding to the isolation of exosomes, the serum samples were first pelleted by centrifugation, and the supernatant was filtered through 0.20-micrometer filtration disk to remove residual cells, debris, platelets, large micro vesicles etc. The samples were incubated with a proprietary precipitation buffer for 1 hour at 4°C. After a 30-minute low speed centrifugation at 500 x g, the exosomes pellets were re-suspended for further analysis of their content. The isolated exosomes was verified by immunoprobing assay using an antibody that recognized exosome surface protein, tumor susceptibility 101 protein (TSG101) (Supplemental Figure-3). Exosome suspension was aliquoted and kept in −20^0^C until use.

### 2.3. Spinal cord homogenate preparation

50-60 milligram thoracic spinal cord tissue (T3 section) was homogenized in an ice-cold buffer (0.32 M sucrose, 0.5 mM MgSO4, 10 mM ε-amino-n-caproic acid, protease inhibitor cocktail (0.05% v/v) (Calbiochem #539134), phosphatase inhibitor cocktails set II (0.1% v/v) (Calbiochem # 524625), 10 mM HEPES, pH 7.4) in a glass pestle-glass homogenizer (8-10 strokes). The tissue/homogenization buffer ratio was one (mg tissue) / 15 (μL buffer). The homogenate was resuspended 8-10 times using a 1 mL syringe with a 26^-gauge^ needle to shear DNA and liberate nuclear proteins. The homogenate was incubated on ice for 20 minutes and subjected to centrifugation at 16,000 X g for 20 minutes at 4^0^C. The supernatant containing all soluble proteins, including nuclear proteins was collected into a clean tube. For purpose of the protein assay, 5-10 μL aliquot was saved. Total protein estimation was analyzed by BCA assay (ThermoFisher Scientific #23225).The sample was aliquoted into 100 μL volumes into microfuge tubes and stored at −80°C until use.

### 2.4.. Immunoblotting analysis

10 μg of exosomal proteins were loaded onto a commercial 4-20% SDS/PAG gradient gel. The electrophoresis was run at 100V for 75-80 min until the dye-front migrated to 0.5 cm from the bottom of the gel. Resolved proteins on the gel were transferred onto a PVDF membrane by electro transfer unit at 60 V for 90 min. The membrane was stained/destained for total protein staining according to manufacturer’s protocol (REVERT Total Protein Stain kit, LI-COR; Cat# 926-11016). The membrane was blocked with a blocking agent (SEA BLOCK Blocking, Thermo Scientific, Cat # UH2788881) for one hour at room temperature, followed by overnight incubation with anti-TDP-43 Ab (1:1000 dilution; Proteintech Cat# 10782-2-AP), anti-P(S409/410) TDP-43 Ab (1:1000 dilution; Proteintech Cat# 66318-1-Ig), and anti-SOD1 Ab (1:750 dilution; Proteintech Cat#10269-1-AP) on an orbital shaker at 4^0^C. The next day the membrane was incubated with Infrared (IR)-tagged antibody (1:10,000 dilution; LI-COR Cat# 926-3211, 926-68071, 926-32210, 926-68070) for 1-2 hours at room temperature. The protein bands were visualized in an image analyser (LI-COR Odyssey Infrared Imager (Model No. 9120). The band intensity was normalized based on total protein staining signals, and analysed by Image Studio image analysing program (V.3.1, Li-COR Biosciences).

## 3. Results

### 3.1. Dogs with DM spinal cord homogenate show elevated TDP-43 and SOD1 in the thoracic region of the spinal cord

The thoracic region of the spinal cord from companion dogs with DM (n=4) and from age matched non-DM affected control dogs (n=4) were obtained from a tissue archive at the University of Missouri College of Veterinary Medicine. The signalment of dogs used in this study listed in **Table-1** in Supplemental Data. **Figure-1** shows that TDP-43 protein levels are elevated in the DM spinal cord region as compared to that of the control dogs. A t-test analysis indicated that differences between two groups were significant (P< 0.01). This observation agrees of other published data that TDP-43 accumulation has been observed in human spinal cord (23, 28). SOD1-immunoreactive aggregates were observed in ventral horn motor neurons (29). This study confirmed that the spinal cord SOD1 aggregation profile occurred in dogs with DM (Fig.2). In this study, the SOD1 aggregation profile was considered as positive control.

**Figure-1.**
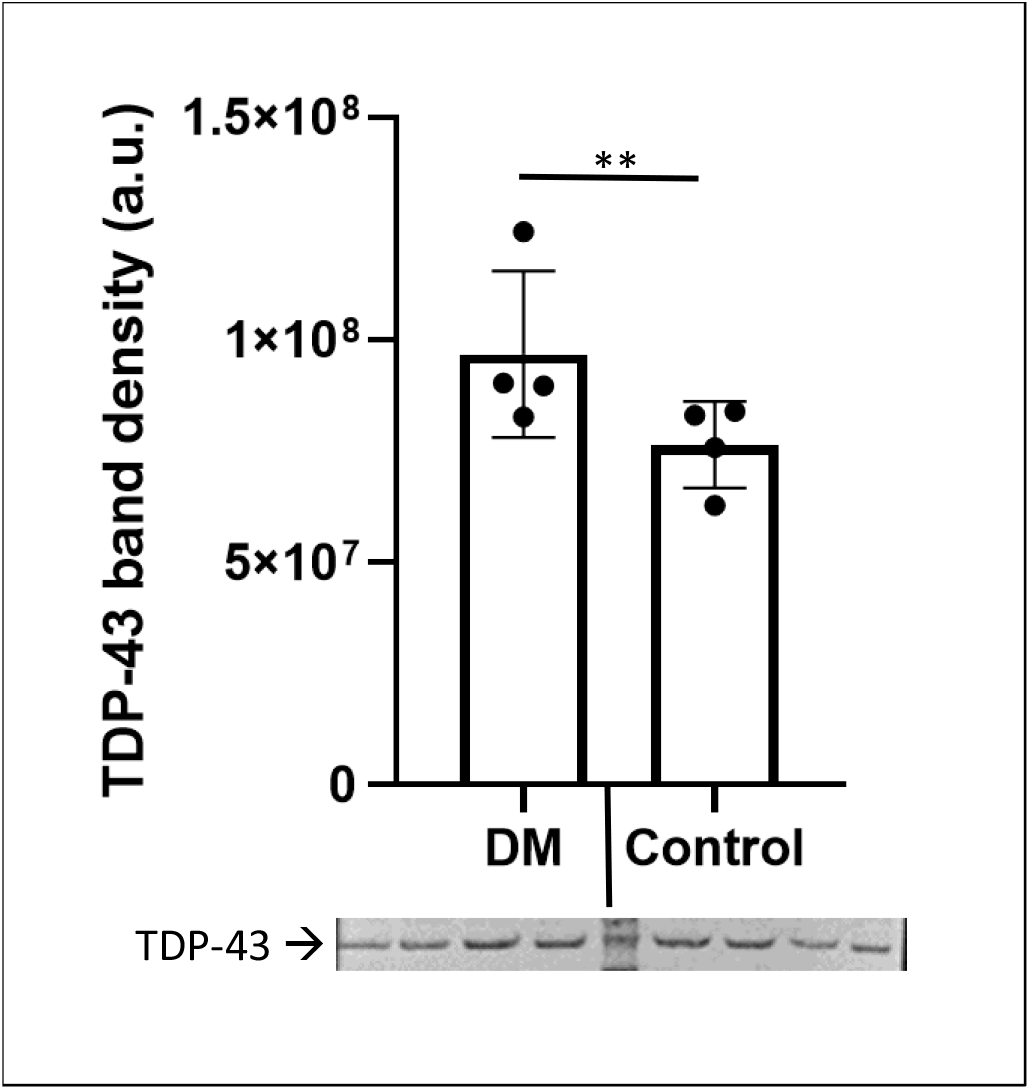
Total TDP-43 levels of thoracic region of spinal cord homogenate. TDP-43 levels were elevated in the spinal cord homogenate from DM affected dogs (n=4, Supplemental Data / Table-1: dog # 1,2,3,4) as compared to that of the control dogs (n=4, Supplemental Data / Table-1: dog # 5,6,7,8) [One-sample t-test (P<0.01)]. The protein band intensity were normalized to total protein staining.

**Figure-2.**
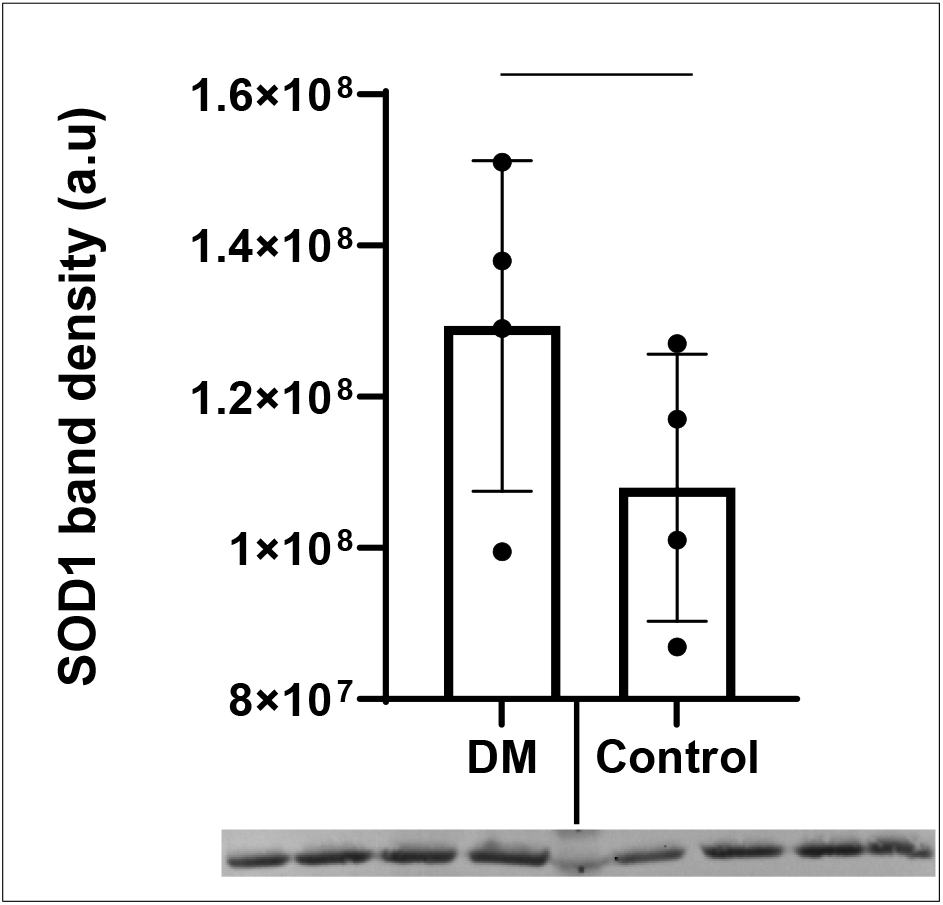
SOD1 levels of thoracic region of spinal cord homogenate. SOD1 levels were elevated in the spinal cord homogenate from DM affected dogs (n=4, Supplemental Data / Table-1: dog # 1,2,3,4) as compare to that of the control dogs (n=4, Supplemental Data / Table-1: dog # 5,6,7,8) [One-sample t-test (P< 0.01)]. The protein band intensities were normalized to total protein staining.

### 3.2. Exosomal TDP-43, phosphorylated TDP-43, and SOD1 are elevated in canine with DM

Frozen serum samples from dogs were obtained from a tissue archive at the University Of Missouri College Of Veterinary Medicine. The signalments of dogs used in this study were listed in **Table-2 and Table-3** in Supplemental Data. **Figure-3 (A, B)** showed that both total TDP-43 and its phosphorylated derivative (pTDP-43) were elevated in exosomes of the dogs affected with DM as compared to that of the control dogs. A significant difference (P≤0.001) was found between DM affect and control dogs. These preliminary results suggest that exosomes can be used to assess biomarker proteins such as TDP-43 and its phosphorylated derivatives.

**Figure 3.**
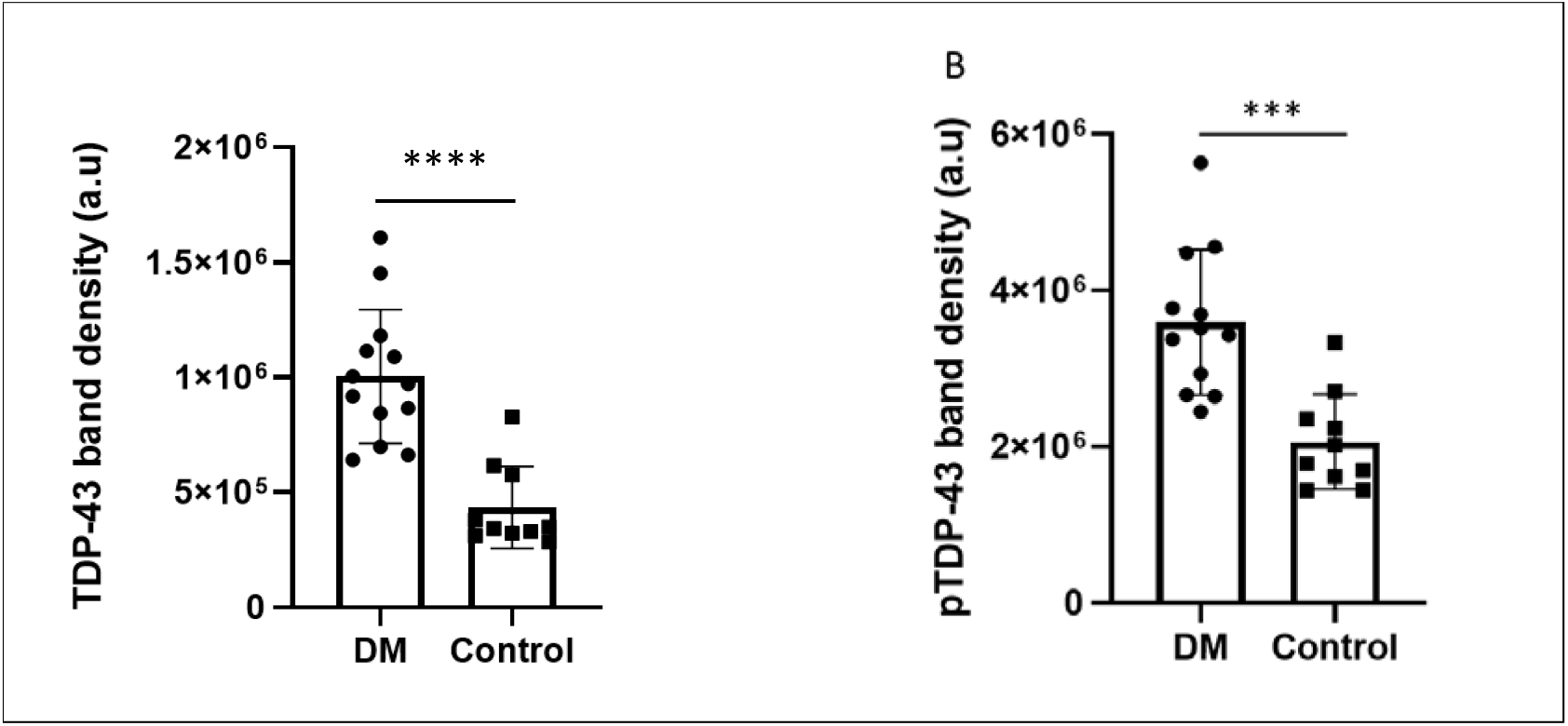
Serum derived exosomal total TDP-43 and phosphorylated TDP-43 profile in DM affected dogs. Both total TDP-43 (A) and phosphorylated TDP-43 (pTDP-43) (B) levels were elevated in serum-derived exosomes from DM affected dogs (n=12; Table-2 Supplemental Data: dog # 1,2,3,4,5,6,7,8,9,10,11,12) compared to control dogs (n=10; Table-3 Supplemental Data: dog# 1,2,3,4,5,6,7,8,9,10). One-sample t-test analysis revealed statistical significance for total TDP-43 (P<0.0001) and for pTDP-43 (P<0.001) Original immunoblot analysis pertinent to this figure provided in Supplemental Fig.1 and Fig.2. The protein band intensities were normalized to total protein staining.

Serum-derived exosomal SOD1 protein levels were elevated in dogs affected with DM as compare to that of control dogs. The difference between the two groups was significant based on an unpaired t-test analysis (P≤0.05). We have observed that SOD1 proteins in canine serum-derived exosomes exhibited a heterogenic profile (i.e., monomer, trimer and tetramer) (Fig.4). Soluble monomers (~25 kDa) were considered in the data analyses and graphing.

**Figure 4.**
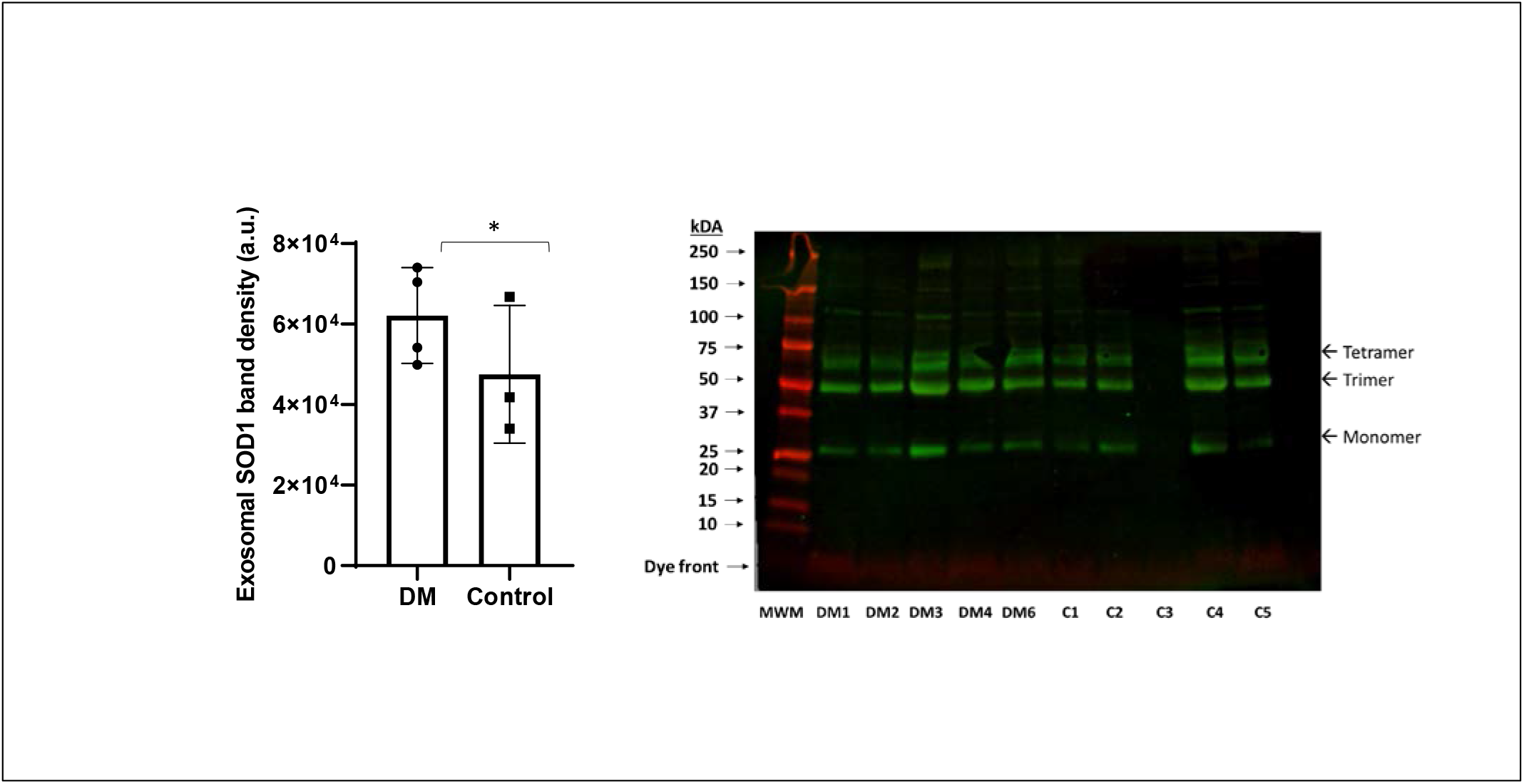
Serum derived exosomal SOD1 profile in dogs with DM; (A) Exosomal SOD1 levels appear to be elevated in dogs with DM. The difference between control (n= 3, Table-3 Supplemental Data; dog #1,2,4,5) and DM (n=4; Table-2 Supplemental Data: dog #1,2,3,4,6) groups did achieve statistically significance (p <0.05). (B) Monomer SOD1 proteins bands were analysed and graphed. The protein band intensities were normalized to total protein staining.

## 4. Discussion

Complexity of a biosample matrix and the invasive nature of tissue sampling led investigators to develop peripheral tissue-based biomarkers for ALS. Blood-based biomarker initiatives have gained popularity due to easy access for sampling with minimal invasion (i.e., venous puncture), access to several blood cells, and subcellular structures. Once developed and validated, blood-based biomarkers will be utilized for diagnosis, monitoring the prognosis, and monitoring the treatment of ALS (30–32).

Canine degenerative myelopathy (DM) is an adult-onset, progressive neurodegenerative disease that shares important similarities to ALS including clinical presentation and progression, pathological features, and *SOD1* mutations (33). This naturally occurring disease in the companion dog recapitulates some forms of human mutant SOD1 ALS. The E40K SOD1 mutation is widespread among companion dogs (34) and leads to SOD1 aggregate accumulations within cells, the putative mechanism of disease in DM and in both familial and sporadic forms of human ALS (16, 35–37).

In this preliminary study, we demonstrated that serum-derived exosomes may be utilized for assessing ALS hallmark proteins (i.e., TDP-43, pTDP-43 and SOD1), and compared to that of thoracic spinal cord homogenate in DM affected dogs. We chose the thoracic region because the disease pathology of canine DM is most severe in this region (33). This region also reflects more TDP-43 accumulation relevant to pathology of ALS (38). Serum exosomes were isolated from the dog cohort consisting of animals with DM and age matched non-DM affected. Due to some technical obstructions and limited tissue amount, we were unable to isolate sufficient exosomes from the spinal cord tissue from this cohort. We are currently refining our techniques to increase the exosome yield.

This study demonstrated the feasibility of the followings: (i) effectively isolating exosomes from serum, (ii) analyse exosomal TDP-43 and pTDP-43 protein profiles, and (iii) comparing those profiles to spinal cord tissue (thoracic region) homogenate from the different cohort of dogs with DM as part of proof-of-concept studies. TDP-43 and SOD1 are known as the hallmark proteins for ALS (23, 24, 39). This study demonstrated that total homogenates of spinal cord TDP-43 levels were elevated in dogs with DM (Fig.1). The SOD1 profile served as a positive control and established that both proteins were elevated (Fig.2). The next logical step would be to demonstrate sensitivity and specificity to establish biomarker reliability within the study limitations due to lack of the biosample archive did not follow the specified minimal information for studies of extracellular vesicles, MISEV2018 (40).

Exosomes are small stable single-membrane organelles that carry selected proteins, lipids, nucleic acids, and glycoconjugates (6). Exosome biogenesis is a valuable mechanism to assess the protein quality control (6). They are emerging as key mediators of communication and waste management among neurons, glial cells and connective tissues during normal and diseases conditions (41). Although exosomes are being developed as therapeutic agents in multiple disease models, they can also serve as surveillance agents by transporting aberrant proteins like hyper-phosphorylated TDP-43 or cytosolic TDP-43 that accumulates in cell plasma. Using dog DM model may establish the link between spinal cord and serum/plasma derived exosomes in terms of TDP-43 biochemistry. We have observed increased TDP-43 and SOD1 levels in exosomes derived from DM affected dog. SOD1 analysis was performed for further confirming the spinal cord and serum exosome crosstalk in terms of the content TDP-43 and its derivative.

The SOD1 protein profile revealed three SOD1 species (monomer, trimer, and tetramer), although SOD1 resolved under the reducing conditions. Recent observations demonstrated that trimer-SOD1 aggressively promoted cell death (42, 43). Because of their small size, exosomes can easily pass the blood brain barrier to serve as accessible biomarkers of neuronal dysfunction (7, 9, 44, 45). This pilot study provided some preliminary data support the notion that spinal cord TDP-43 may be transported via exosomes, and that TDP-43 loaded exosomes appear in blood. However, we did not perform spinal cord-derived exosomal TDP-43 and SOD1 assessment. Hence, we do not claim yet serum-derived exosomal TDP-43 and SOD1 directly reflecting that of spinal cord. Studies are undergoing to further isolate spinal cord derived exosomes appeared in serum and analyze their cargo content in terms of TDP-43 and SOD1.

The concept of serum-derived exosome analysis for potential biomarker candidates will eliminate very invasive tissue and cerebrospinal fluid sampling and allows investigators to obtain repeated blood samples with a minimal invasive sampling method for exosome isolation. Exosomes derived from serum represent global exosomes which include neuronal derived exosomes (NDE) (46). A recent review has discussed on astrocyte-derived extracellular vesicles in the blood may be a useful tool for detecting astrocyte-specific biomarkers in neurological conditions (47, 48) Our current research is focusing on the development of a process, to selectively isolate the NDE population from global serum and spinal cord tissue derived exosomes.

The availability of a disease model that more closely reflects the human condition would provide an opportunity to test if the pathologic processes shown to contribute to the motor neuron disease in rodent models are likely to contribute to ALS. Such a model could also be used in second-generation preclinical trials to shield ALS patients from participation in clinical trials with little probability of success. In addition, studies with the proposed disease model may reveal disease processes that are shared with the ALS cases but do not occur, or are difficult to detect, in the rodents due to small size, short lifespan, or other issues. Ideally, the desired disease model would develop both upper and lower motor neuron signs due to a single spontaneous mutation in an orthologue of a human gene known to harbor ALS-causing mutations. This model would involve an animal species that is intermediate between rodents and humans in life expectancy and in the size and complexity of the CNS. It would be readily available on a wide variety of genetic backgrounds negating the need to maintain research colonies. It would display a disease phenotype that, similar to ALS, first manifests in mature individuals. In addition, the initial clinical signs would consistently appear at the same anatomic site and spread throughout the body in a uniform in pattern and rate. Canine DM has all these features and is therefore another disease model for understanding ALS. Companion dogs with DM represent a clinical population confounded by complexities in diagnosis, comorbidity and environmental and genetic diversity similar to those encountered in a human clinical trial setting. Thus, incorporation of veterinary clinical trials into the ALS treatment development paradigm will enhance translational efficiency by identifying and optimizing those therapies most likely to generate clinical benefit.

## Acknowledgments

Authors acknowledge the Office for Research and Support Program at Kansas City University for supporting student research and providing logistic supports, Robert Liu, DO and Qwynton Johnson for providing technical help, Mrs. Jessica Sage for training summer research fellows, and Dr. Jan Talley and Ms. Tuba Agbas for reading and editing the manuscript.

## Author Contributions

Conceptualization, AA and JRC; methodology, AA, PP, AK,KG, YME, MM and EK.; formal analysis, PP,AK,KG,EK, and AA; writing, original draft preparation, AA; writing, review and editing, AA,MM, and JRC; supervision, AA and EK; All authors have read and agreed to the published version of the manuscript.

## Data availability statement

The authors confirm that the data supporting the findings of this study are available within the article and its supplementary materials. The data are available on reasonable request from corresponding author

