## Supplemental figures for "Exosomal TAR DNA binding protein 43 profile in canine model of amyotrophic lateral sclerosis: A preliminary study in developing blood-based biomarker for neurodegenerative diseases"

### Raw data for Figure-1 and Figure -2

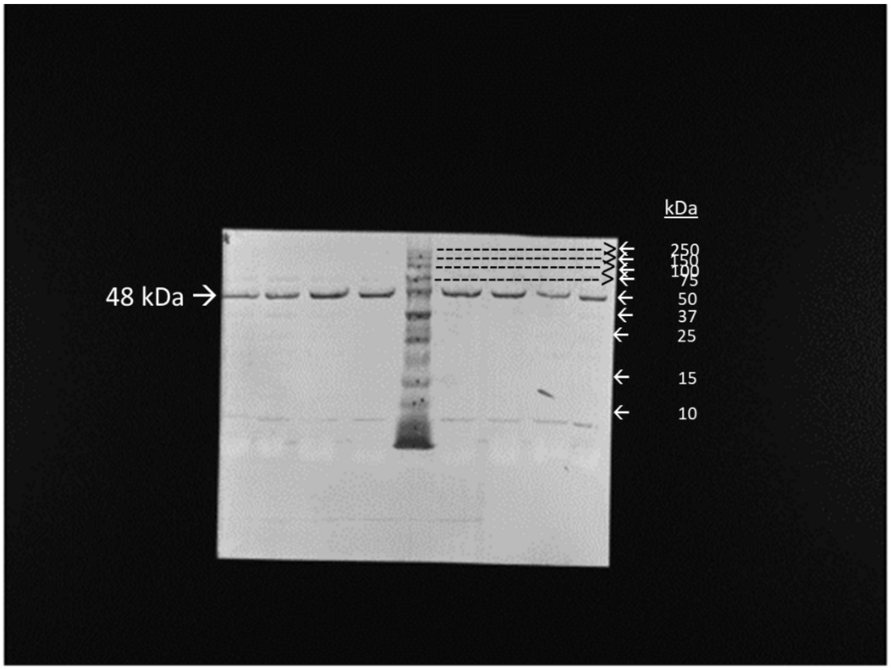


Original uncropped image for TDP-43 western blot in Figure-1

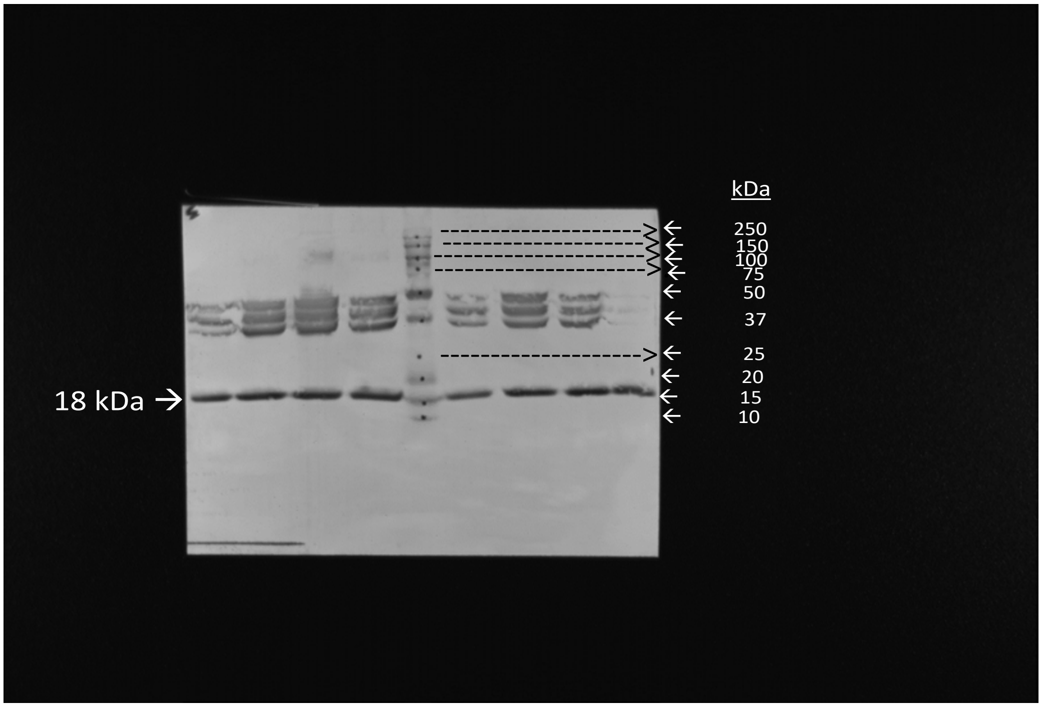


Original uncropped image for SOD1 western blot in Figure-2

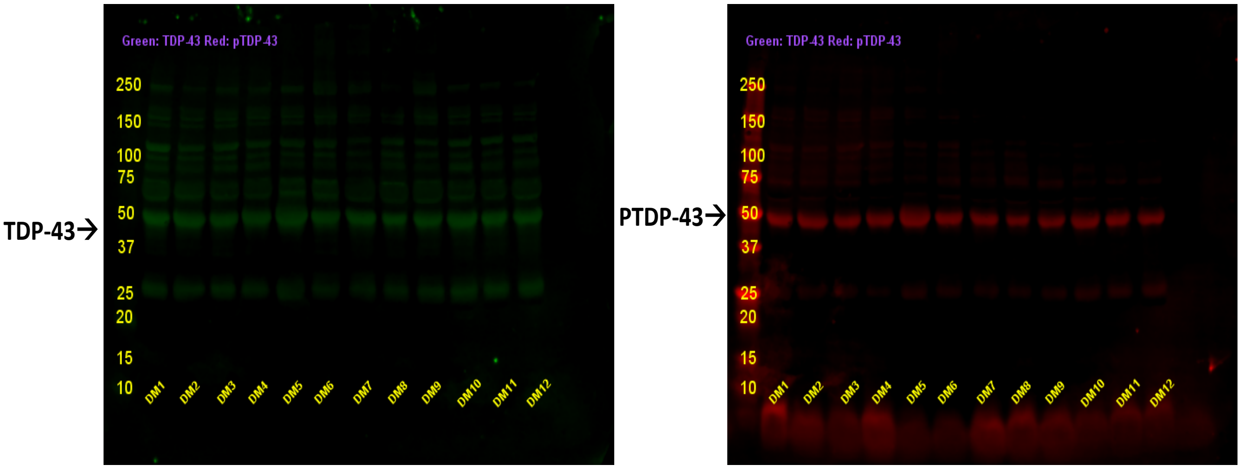


**Supplemental Figure-1.** Serum-derived exosomal TDP-43 and phosphorylated TDP-43 profile in dogs with DM (n=12; Table-2 Supplemental Data : dog # 1,2,3,4,5,6,7,8,9,10,11,12)

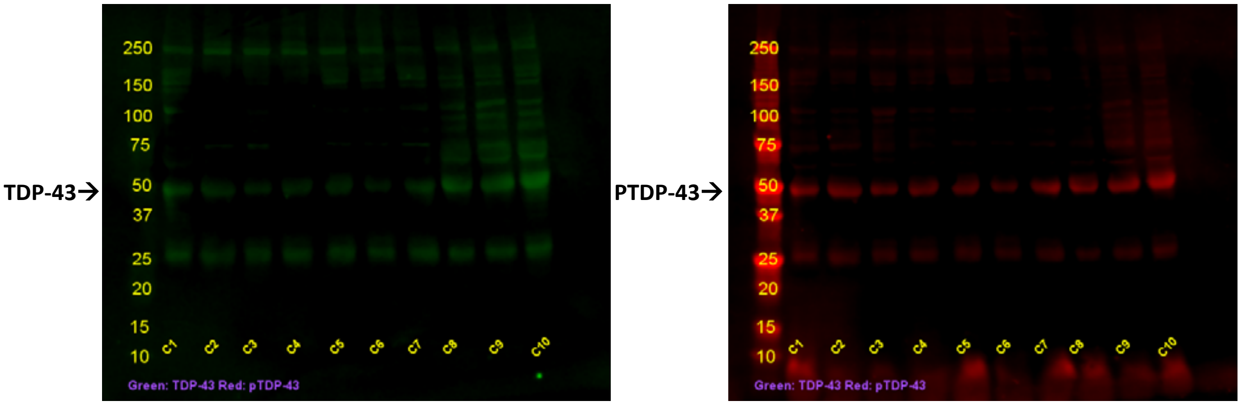


**Supplemental Figure-2 .** Serum derived exosomal TDP-43 and Phosphorylated TDP-43 profile in control dogs (n=10; Table-3 Supplemental Data : dog # 1,2,3,4,5,6,7,8,9,10 )

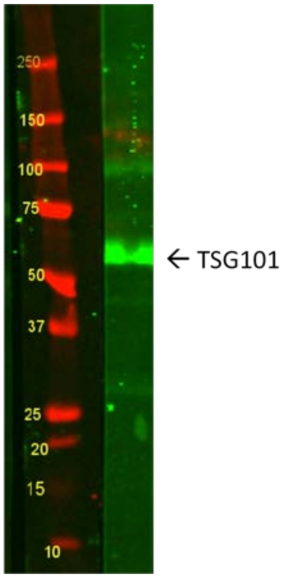


**Supplemental Figure-3**. Serum/plasma derived exosome isolation and verification by a-TSG101 antibody.

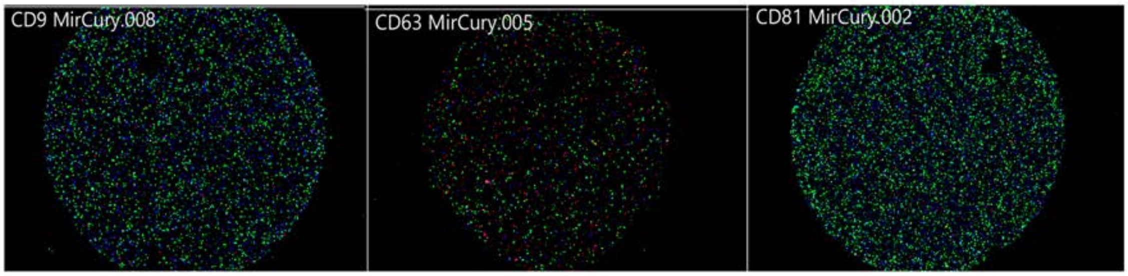


**Supplemental Figure-4** : Serum-derived exosome imaging by ExoView platform. The images show the binding of single exosomes to CD9 (Blue), CD63 (Red), and CD 81(Green) antibody spots. Serum/plasma derived exomes obtained by miCURY exosome isolation kit (QIAGEN) were analyzed by ExoView platform for their population.
